# Membrane channel hypothesis of lysosomal permeabilization by beta-amyloid

**DOI:** 10.1101/2021.07.31.454606

**Authors:** Dmitry V. Zaretsky, Maria V. Zaretskaia, Yaroslav I. Molkov

## Abstract

Alzheimer’s disease (AD) is the most common cause of dementia affecting millions of people. Neuronal death in AD is initiated by oligomeric amyloid-β (Aβ) peptides. Recently, we proposed the amyloid degradation toxicity hypothesis, which explains multiple major observations associated with AD – such as autophagy failure and a decreased metabolism. According to the hypothesis, the key event in the cellular toxicity of amyloid is the formation of non-selective membrane channels in lysosomal membranes by amyloid fragments that are produced by the digestion of Aβ previously absorbed by endocytosis. Electrophysiological data suggest that amyloid-formed channels have different sizes, which can be explained by the fact that barrel-shaped amyloid aggregates which create channels can consist of different number of monomers.

To estimate the ability of channels to leak molecules of various molecular weights, we modeled the channels as saline-filled cylinders in non-conductive membranes that pass spheres with a density of average globular proteins. As a basis, we used the conductance distribution taken from the previously published experimental dataset, in which single channels with a conductance reaching one nanosiemens were registered. Our calculations show that channels with a giant conductance can allow for passing macromolecules such as lysosomal cathepsins implicated in the activation of apoptosis. The formation of giant channels is disproportionally promoted in an acidic environment. Also, amyloid fragments leaking from permeabilized lysosomes can reach the internal leaflet of the plasma membrane and permeabilize it.

We conclude that while dissipation of the proton gradient by any – even the smallest amyloid channel – readily explains lysosomal failure, the relatively rare events of lysosomal permeabilization to large macromolecules can be an alternative mechanism of cellular death induced by exposure to Aβ.

## Amyloid channel vs Amyloid degradation toxicity hypothesis of Alzheimer’s disease (AD)

The mechanisms of Alzheimer’s disease have been under intense investigation in recent decades, but still, no integrative theory of AD exists. Senile plaques that consist of beta-amyloid peptides (Aβ) are the best-known anatomical markers of the disease, but they are not considered the cause of the disease. The cytotoxicity of Aβ is mediated by oligomers, while monomers and fibrils are relatively non-toxic [12].

About thirty years ago, several groups independently reported that Aβ forms non-selective membrane channels with an electrical conductance of up to several nanosiemens [4-6, 21-23, 25]. The concept that these channels are barrel-shaped structures formed by multiple peptide molecules [10] supports the role of amyloid oligomers in Aβ toxicity [9, 27]. However, there are multiple features of AD that are not explainable by the amyloid ion channel theory in its original form [37]. One of them is lysosomal failure and autophagy disfunction. Autophagic vacuoles (AV) are rare in the neurons of the normal adult brain, but their presence is dramatically increased in Alzheimer’s disease [24] and is most likely caused by a failure of AV maturation [24]. Immunolabeling identifies AVs as a major reservoir of accumulated intracellular Aβ [34].

Aβ is internalized through endocytosis [15, 18, 30]. Intracellular fluorescently labeled Aβ creates a punctate pattern of fluorescence and shows a strong correlation with the lysosomal marker (Fig.1A, B). Similarly, the cells accumulate the membrane-impermeant Lucifer Yellow (LY, MW 444), which enters the cells through endocytosis [17]. In untreated cells, LY was observed in the form of circumscribed vesicular structures. In contrast, after Aβ_1–42_ treatment for 24 h, the cells displayed a diffuse pattern of fluorescence, suggesting that lysosomal content leaked to the cytoplasm in a significant percentage of cells. Exposure to a high concentration of Aβ resulted in cellular degeneration, which is manifested by the rounding of the otherwise flat attached cells. However, the leakage also occurred in some cells that are not rounded, so lysosomal permeabilization precedes the manifestation of cellular degeneration.

**Figure 1.**
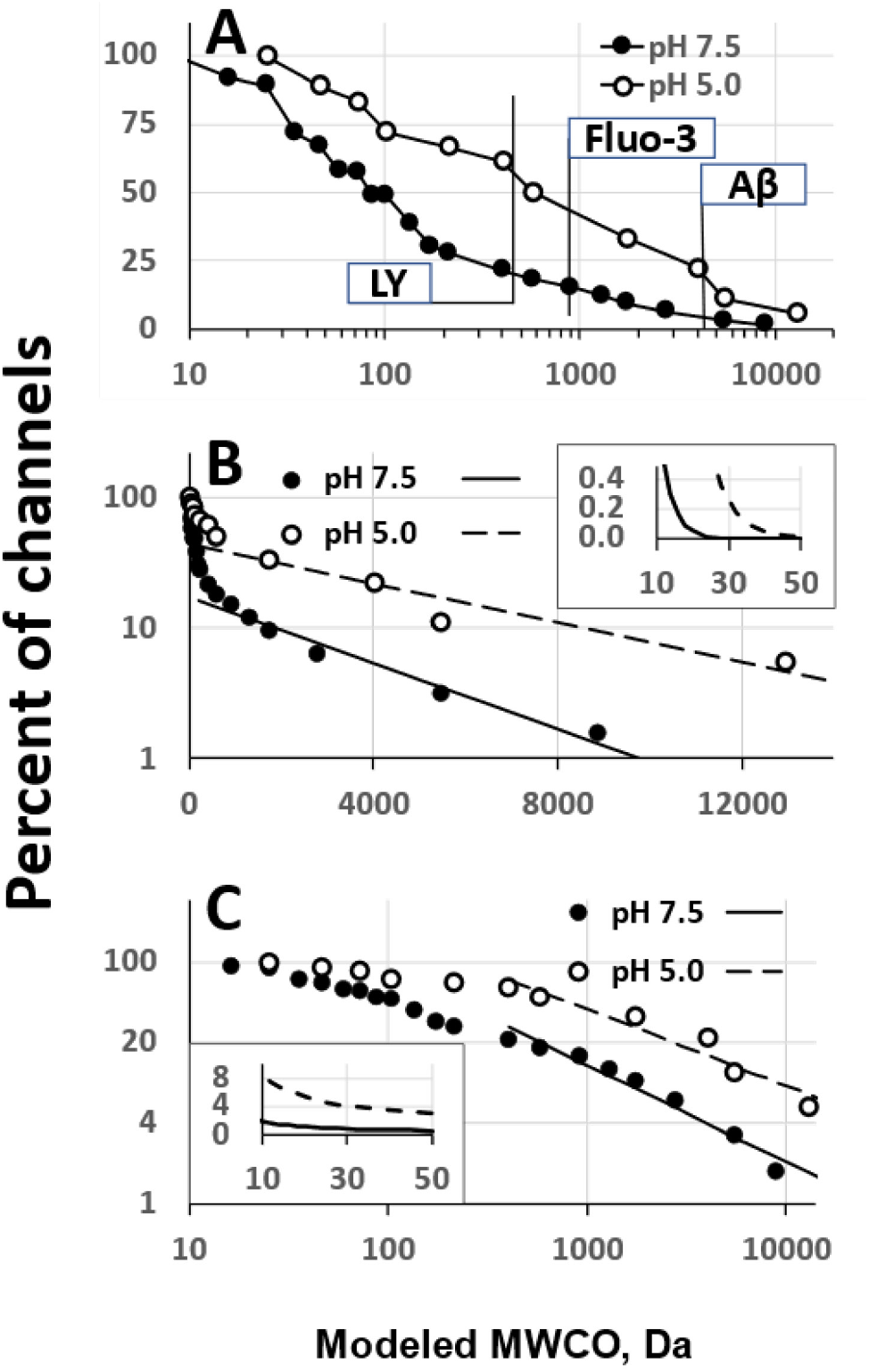
The percentage of channels with a conductance which in model conditions is sufficient to pass globular proteins of various molecular mass. The datasets on conductance distribution of channels created by Aβ_25-35_ at pH 7.5 and 5.0 are taken from Lin&Kagan, 2002 [22]. Two curves are different if compared by Kolmogorov-Smirnov test (p<0.05). The dependence is presented in different coordinates to estimate the way to extrapolate the probability of giant channel formation. Linear fitting of distributions was performed for conductances which correspond to MWCO exceeding 500 Da. **A**. Log-linear scale of percentage vs MWCO to show the overall distribution and molecular weights of some relevant compounds - Lucifer Yellow (LY), calcium-sensing probe Fluo-3 and monomeric Aβ_1-42_. **B**. Linear-log scale to estimate exponential dependence. **Insert:** Extrapolation of probability of giant channels based on exponential fitting for MW up to 50 kDa. **C**. Log-log scale to estimate dependence as a power function. **Insert:** Extrapolation of probability of giant channels based on fitting with a power function for MW up to 50 kDa.

Aβ-induced lysosomal permeabilization is well described [17, 30, 32]. However, the molecular mechanism of this phenomenon was never conclusively linked to a specific biological substrate. Unlike the abnormally slow processing of autophagosomes, which is a functional deficiency, the permeabilization of lysosomes can be viewed as a breach of their structural integrity. Lysosomes can leak molecules much larger than LY. The appearance of lysosomal enzymes in the cytoplasm also supports the notion that the cytoplasmic spread of LY in amyloid-treated cells does not result from the permeabilization of plasma membranes [17].

In connecting lysosomal permeabilization with amyloid channel formation, we considered two key facts. First, except for the areas of lipid rafts that are enriched in gangliosides, the external leaflet of plasma membranes does not carry a significant negative surface charge, which is necessary for the channel incorporation [3, 36]. In contrast, lysosomal membranes contain a significant ratio of negatively charged bis(monoacylglycero)phosphate [2]. Second, a full-length peptide, i.e. Aβ_1–42_, is not efficient in creating membrane channels, unlike its fragments – such as Aβ_25–35_ [23, 35]. Along with being a target for membrane channel formation, lysosomes provide proteases which can produce channel-forming amyloid fragments. The digestion of Aβ, producing channel-forming fragments, is the etiological factor of AD in the amyloid degradation toxicity hypothesis [37].

The goal of this study is to theoretically discuss the following questions: can membrane channels be a molecular substrate of lysosomal permeabilization induced by beta-amyloid, and what is the maximal size of the molecules the lysosomes can become permeable to? To answer these questions, we used a mathematical model to estimate how capable amyloid channels are in passing molecules of various molecular weights based on electrophysiological data published earlier in [22].

## METHODS

### Conductance of membrane channels formed by Aβ_25-35_

The dataset for the analysis was taken from the study by Lin et al. [22]. In this study, black lipid membranes consisting of various lipids were formed over small holes (200 or 1000 µm in diameter). The membranes were exposed to solutions of amyloid fragment, and a current passing through created channels was recorded in voltage clamp conditions. Experiments were performed at various pH in the range of 4.5–9.9. A histogram of single channel conductances was constructed by observing all the current jumps, including the initial insertion into the membrane and subsequent openings of channels.

### Mathematical modeling of single-channel conductance as a measure of molecular weight cut-off (MWCO)

To estimate the size of the proteins that can pass through open channels, we calculated the molecular weights of spheres with the density of globular proteins that can pass through channels with various conductances corresponding to the conductances of amyloid channels described in [22]. The membrane was considered non-conductive, while membrane channel was modeled as a cylindrical pore filled with saline that has resistivity *p* (Ohm · cm). The length of the pore *l* (cm) is equal to the thickness of membrane. The diameter of such a pore *D* (cm) can be calculated as a function of conductance *g* (Ohm^−1^) using the formula [7]:

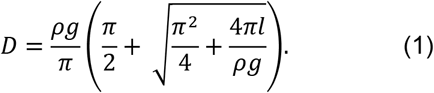

We assumed that the protein has the simplest shape, a sphere, and an average density of globular proteins of 1.37 g/cm^3^ [11]. Then, the radius of the protein molecule *R* (nm) can be calculated as follows:

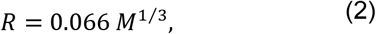

where *M* is the molecular weight in Da. Using (1) and (2) and that *D* (cm) = 2 · 10^−7^*R* (nm), the molecular weight of the largest protein passing through the channel (molecular weight cut-off, MWCO) can be linked to the conductance by

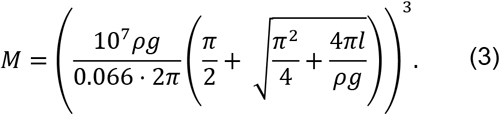

Following the approach of Bode et al., for calculations we used the thickness of membranes *l* = 7 nm and that the solution resistivity *p* = 80 Ohm · cm [7]. To express *g* in pS and *l* in nm, we recalculated

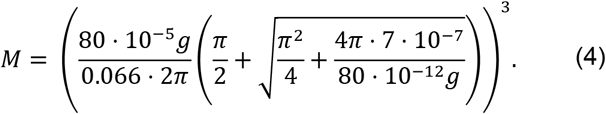

After simplification, the formula is

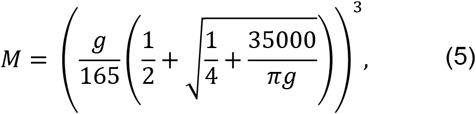

where *M* is the MWCO in Da and *g* is the channel conductance in pS.

## RESULTS

### Electrophysiological data on conductance of membrane channels formed by Aβ_25-35_

As shown by Lin and Kagan [22], most membrane channels formed by this 11-amino acid peptide had conductances between 10 and 200 pS, with some reaching 1 nS. The acidification of the solution did not change the number of formed channels [22], but shifted the distribution towards significantly higher conductance values. The molecular weights of protein molecules, which are theoretically passable through the membrane openings with corresponding conductances were calculated as described in Methods.

After recalculating channel conductances into MWCO, we constructed the distribution of channels by MWCO as shown in log-linear coordinates in Fig.1A. The distribution of channels at pH 5.0 appears significantly different from the one at pH 7.5. The molecular weights of several molecules relevant to the discussion are shown in Fig.1A: Lucifer Yellow (LY, MW 444), Fluo-3 (MW 770), and monomer of Aβ_1-42_ (MW 4514).

### The probability of channels with high MWCO

The probabilities of channels with high MWCO values that were not registered experimentally were extrapolated using assumptions about asymptotic behavior of the distributions. We tested two options: when the channel occurrence probability is an exponential or a power function of MWCO.

To extrapolate the probability distributions under the first assumption, the data was plotted in linear-log coordinates and linearized by fitting the data points corresponding to MW>500 Da (Fig.1B). Extrapolation of these lines to higher values demonstrate that there is a significant occurrence probability of channels able to pass macromolecules with a molecular weight exceeding 10 kDa (Fig.1B, insert). This probability exponentially decreases with size based on the underlying assumption. In an acidic environment, the probability curve appears shifted by about 20 kDa compared to neutral conditions and apparently has a slower decay.

The asymptotic approximation of the distributions by a power function provides even higher estimates of giant channels. To perform these extrapolations, we represented the data in log-log coordinates, and, again, found a linear fit of data points with MW>500 Da (Fig.1C). Extrapolation to the MWCO>10 kDa expectedly provides much higher probabilities for giant channel occurrence compared to the exponential approximation (Fig.1C, insert). Even at pH 7.5, the ratio of channels passing 50 kDa molecules is approximately 0.5% of the total number of all formed channels. At pH 5.0, which promotes the formation of channels with a higher conductance, this probability is almost 3% and remains above 1% for the MWCO>200 kDa (not shown).

## DISCUSSION

### Lysosomal membranes are a probable target for beta-amyloid

The amyloid channel hypothesis remains one of the major theories of beta-amyloid cytotoxicity since the phenomenon of amyloid channel formation was discovered three decades ago in artificial lipid membranes. Unfortunately, in its current form, this hypothesis falls short of explaining multiple aspects of amyloid toxicity, as well as provides a very limited insight into the pathophysiology of AD.

The typical assumption of the amyloid channel theory that pathophysiological consequences of amyloid exposure result from its effects on the plasma membrane does not seem to hold and has been distracting the researchers’ attention from the potential roles of other cellular compartments that may also be affected. Lysosomes are a good research target, as their membranes carry a negative surface charge due to a significant ratio of bis(monoacylglycero)-phosphate [2]. Unlike other negatively charged cellular membranes – e.g. the inner membrane of mitochondria – after the cell is exposed to Aβ, lysosomal membranes quickly become exposed to the amyloid because the amyloid is up-taken by endocytosis [15, 18, 30]. Furthermore, lysosomes are unique in the cellular machinery due to their role in protein degradation. The production of amyloid fragments provides the materials that are able to effectively form membrane channels, unlike full-length Aβ. Taken together, the metabolic function of lysosomes and the negative charge of the lysosomal membrane make these organelles a primary target for the formation of channels.

Recently proposed by us “amyloid degradation toxicity hypothesis” of AD etiology and pathogenesis builds on these arguments and proposes that channel formation in lysosomal membranes by amyloid fragments is an initial molecular insult, initiating a biochemical pathway leading to cell disfunction and eventual cell death [37].

### Amyloid channel formation provides a biologically relevant mechanism for lysosomal permeabilization

The exposure of a cell in the culture to exogenous Aβ results in the accumulation of peptide inside endocytic vesicles, which fuse with lysosomes. Using fluorescently labeled monomeric peptides (1 µM), it was estimated that over a period of 8 h, each cell accumulates about 800,000 molecules of Aβ_1–42_ almost exclusively associated with vesicles labeled by a lysosomal tracker [30], and after 24 h retains around 40% of the peptide.

It is important to consider that Lysotracker identifies lysosomes by their acidic content. However, at first, amyloid is enclosed in endosomes, which become progressively acidic during maturation [16] or after merging with lysosomes [14, 33]. Therefore, freshly internalized peptide can be in the vesicles that are not acidic yet, which can explain why some internalized peptide is not co-labeled with Lysotracker. Also, some lysosomes can lose acidity due to the permeabilization, and, therefore, are not labeled with Lysotracker.

The leakage of lysosomal marker Lucifer Yellow into the cytoplasm after the cells are exposed to exogenous Aβ [17] implies that along with small amyloid channels permeable for ions, there are larger channels that can pass larger molecules [26]. The percentage of such larger channels is relatively small, so most of vesicles with channels remain impermeable to small compounds like calcium-sensing probe Fluo-3, which makes flow cytometry useful for studying the channels [36]. Unlike Lucifer Yellow, the internalized fluorescent Aβ itself was observed as a punctate pattern without clear diffusion into the cytoplasm. The absence of clear leakage of Aβ could be due to higher MW compared with LY. A significant percentage of channels are large enough to leak small compounds like LY (Fig. 1A), but only a small fraction of channels can pass large molecules with MW>4000. Also, the concentrating of Aβ up to 100-fold intralysosomally [30] promotes aggregation, so the effective MW of the peptide can be greater than 4 kDa.

The extrapolation of experimentally measured distributions of channels by size predicts that very large channels are rare, but they still are likely to be formed (Fig.1). According to the model that describes the asymptotic behavior of the size distribution by a power function, there is a finite occurrence probability for the channels able to pass molecules larger than 50 kDa (Fig.1C). This prediction is supported by the observation that lysosomal enzyme β-hexoseamini-dase (MW 150 kDa) can be detected in the cytoplasm after exposure to cytotoxic concentrations of Aβ [17].

In this context, lysosomal proteolytic enzymes are not extremely large – cathepsins have a MW of less than 50 kDa, with most of them having MW in 20-30 kDa range. Both distribution models (with exponential and power tails) predict the occurrence of the channels capable of passing molecules of these sizes (Fig.1B, C). If these enzymes leak, and the anti-proteolytic defense in the form of cytoplasmic cathepsin inhibitors fails, degradation enzymes produce cellular necrosis by digesting the cellular content. Alternatively, by activating caspases, they initiate apoptotic mechanisms also leading to cell death.

Clearly, more focused additional studies are needed to characterize the actual processes defining the permeability of amyloid channels to various molecules. A major limitation of our speculations is that our channel permeability model is based on unrealistic assumptions, such as a spherical shape of the protein, an ideal cylindrical shape of the pore, and the ability of the channel to pass a molecule if the diameter of the macromolecule does not exceed the diameter of the pore. Direct measurements could reveal the actual permeability of membranes induced by amyloid channel formation. We predict that despite extremely large channels being rare, they can still occur in physiologically relevant quantities.

### Lysosomal permeabilization is a plausible mechanism of amyloid-induced cellular death

As noted, the non-selectivity of amyloid channels to cations was discussed almost exclusively with respect to the ions of metals. Surprisingly, we could not find any direct studies demonstrating that amyloid channels can pass protons. For example, observed oscillations of pH after amyloid exposure were interpreted as a consequence of Ca^2+^/H^+^ exchange after calcium homeostasis was perturbed [1]. However, it is straightforward to assume that amyloid channels are permeable to H^+^ due to non-specificity of amyloid-induced membrane permeabilization, which would explain why fluxes of calcium and pH were synchronous in cells exposed to Aβ [1]. Indeed, amyloid channels pass all the cations studied – sodium, potassium, calcium, cesium, and lithium [6].

The permeability of amyloid channels in lysosomal membranes to protons readily explains lysosomal/ autophagy failure. Lysosomal proteases are mostly acidic hydrolases and require the actively supported acidification of the lysosomal lumen, so if intra-lysosomal pH equilibrates with the cytoplasm due to channel formation, the digestion process stops. The lysosome then becomes inactive and requires utilization [37]. In the amyloid degradation toxicity hypothesis, the consequences of lysosomal permeabilization initiated by channel formation represent the pathophysiological pathways

The fact that amyloid channels can have different sizes may be the key to understanding the pathophysiology of Alzheimer’s disease. The initial event is amyloid entry through endocytosis. After endocytosed Aβ is degraded, some fragments can form channels, and due to the properties of the lysosomal membrane, can incorporate into the lysosomal membrane. The size of the formed channels defines the consequences of the lysosomal permeabilization. These consequences are summarized as three possible paths depending on the MWCO of the formed channel (Fig.2).

**Figure 2.**
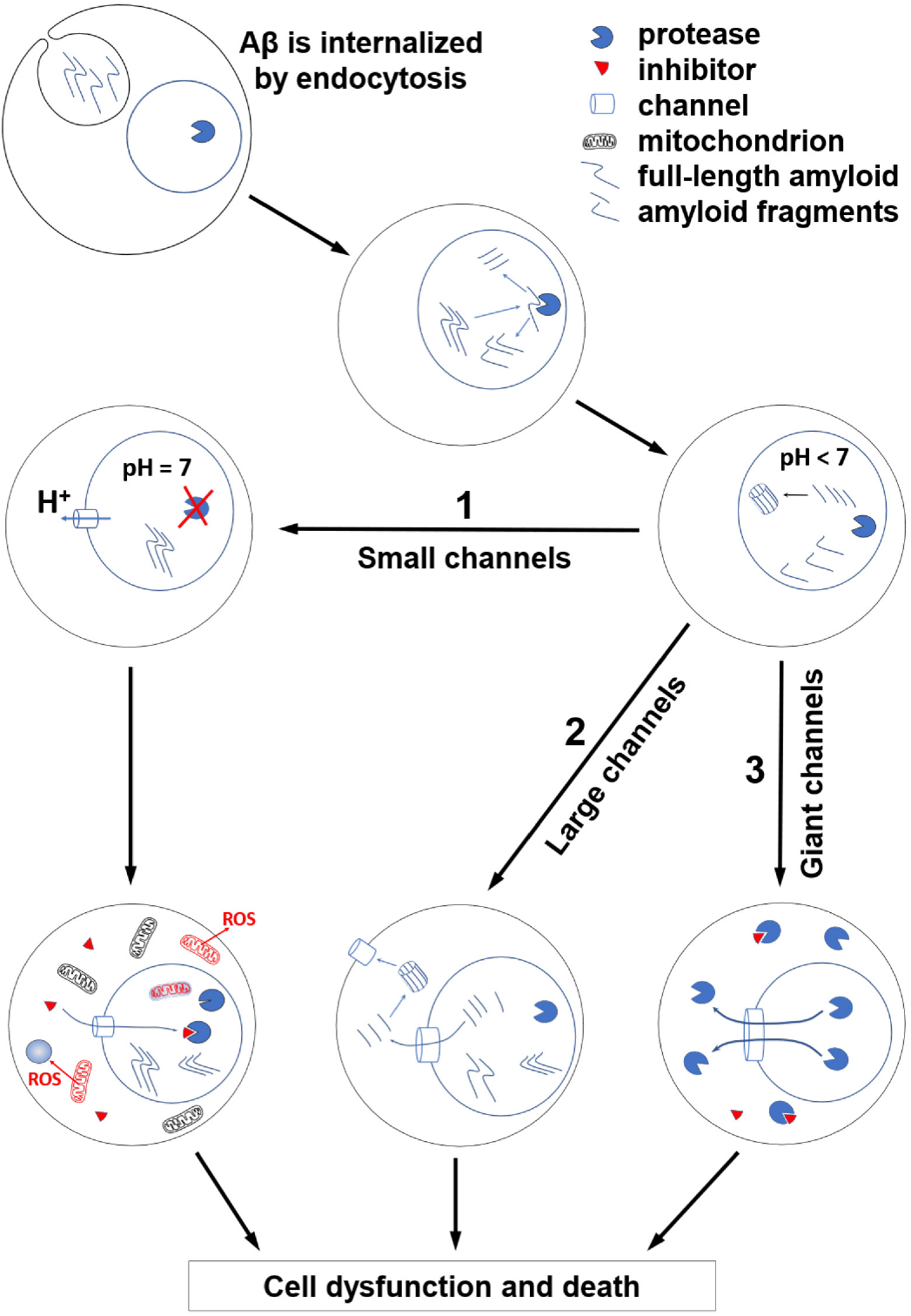
Lysosomal permeabilization and cell death. The endocytic vesicle containing the amyloid peptide is merged with a lysosome. Proteases produce various short fragments and some of them can form non-selective membrane channels. There are at least three pathophysiological pathways resulting in cellular dysfunction and death initiated by permeability of lysosomal membrane. **1. Small channels** (MWCO<200 Da) dissipate the pH gradient. The neutralization of lysosomal lumen inhibits acidic proteases. Dysfunctional lysosomes accumulate, and the recycling of organelles fails. Damaged mitochondria are not recycled and produce reactive oxygen species, damaging other organelles. If channels are large enough, cytosolic protease inhibitors can enter the lysosomes and suppress proteolytic degradation further. **2. Large channels** (MWCO>4 kDa) leak amyloid fragments to the cytoplasm, where channel-forming fragments target other membranes, including the plasma membrane. Plasma membrane loses ionic gradients affecting the ability of neurons to generate action potentials even before the cell eventually dies. **3. Giant channels** (MWCO>10 kDa) leak lysosomal enzymes to the cytoplasm. Some of leaked proteases are inactivated by intracellular inhibitors (cystatins, serpins etc). However, if the leakage is overwhelming, inhibitors cannot protect, and proteases digest the cellular content causing necrosis. Also, leaking cathepsins can activate caspases and initiate apoptosis.

1. All channels, even the smallest ones, can pass protons, thus leading to autophagosomal dysfunction as explained above (Fig.2, pathway 1). In experimental models, the suppression of proteolysis by pharmacological deacidification of lysosomes slows their axonal transport and causes their selective accumulation within dystrophic axonal swellings [20]. Therefore, it is reasonable to suggest that amyloid channel-mediated lysosomal dysfunction also results in disrupted vesicular traffic, which is another prominent feature of AD.
2. Amyloid fragments can leak through larger channels formed in the lysosomal membranes (Fig.2, pathway 2), since the molecular weight of undecapeptide Aβ_25-35_ is above 1 kDa. As shown in Fig. 1, the channels with a MWCO sufficient to pass such peptides (MW 1-2 kDa) and even their oligomers can represent a significant fraction of the formed channels. The leaked fragments would be able to incorporate into the plasma membrane from the side of the inner leaflet, which, unlike the outer leaflet, carries negatively charged phosphor-lipids necessary for the channel-membrane interaction [35]. This hypothesis answers the question of why the exposure to beta-amyloid causes permeabilization of the plasma membrane in living cells with a significant delay of 5-15 min [1], compared to artificial liposomes where channel formation is virtually immediate [36].
3. Another pathway linking lysosomal permeabilization with cell death involves the leakage of lysosomal enzymes (Fig.2, pathway 3). Cathepsins (MW 20-50 kDa) are unlikely to pass through most amyloid channels. This is why in our modeling we paid special attention to the extrapolation of probabilities of large channels in experimental datasets into the range corresponding to these giant channels. In the cytoplasm, lysosomal enzymes either directly digest cellular content (necrosis) [31] or activate cytoplasmic caspases to induce apoptosis-mediated cell death [13, 19, 29]. Despite an acidic pH level (actively supported in the lysosomal lumen) being critical for the optimal activity of proteolytic enzymes, released proteases can remain active in the neutral pH of cytosol for at least some time [28]. Cytosol contains inhibitors that guard against the undesirable leakage of enzymes [8], but this protection can fail [13].

From the early studies of amyloid channels, it was known that blockers are more efficient against membrane permeabilization caused by smaller channels [6]. If cellular death is the consequence of enzymatic leakage, which requires giant channels, AD progression cannot be affected by typical amyloid channel blockers with a low molecular weight such as tromethamine, even though such inhibitors can be effective *in vitro* [6]. This suggests that measuring the MWCO distribution of the amyloid channels experimentally, especially in the range of 10-50 kDa, is one of the important objectives for future experiments.

## CONCLUSION

For over thirty years, the amyloid channel theory claimed membrane channels to be a primary molecular mechanism of beta-amyloid toxicity. However, the traditional concept of plasma membrane channels causing the disturbance of ion gradients falls short when interpreting AD pathophysiology and cell death associated with exposure to amyloid. In this study, we reprocessed previously published electrophysiological data on amyloid channels formation. Based on the results, we argue that beta-amyloid toxicity may be associated with lysosomal permeabilization via the formation of membrane channels with a wide range of molecular weight cut offs. This explains a variety of AD features, including low brain metabolism, autophagy failure, and cell death.

## Acknowledgements

The authors thank Daniel Zaretsky for his editorial help. Research reported in this publication did not receive external funding.

## Abbreviations

AD: Alzheimer’s disease
Aβ: beta-amyloid
MW: molecular weight
MWCO: molecular weight cut-off

## REFERENCES

[1] A.Y. Abramov, L. Canevari, M.R. Duchen, Calcium signals induced by amyloid β peptide and their consequences in neurons and astrocytes in culture, Biochimica et Biophysica Acta (BBA) - Molecular Cell Research 1742 (2004) 81–87.

[2] Z. Akgoc, M. Sena-Esteves, D.R. Martin, X. Han, A. d’Azzo, T.N. Seyfried, Bis(monoacylglycero)phosphate: a secondary storage lipid in the gangliosidoses, Journal of lipid research 56 (2015) 1006–1013.

[3] J.M. Alarcon, J.A. Brito, T. Hermosilla, I. Atwater Mears, E. Rojas, Ion channel formation by Alzheimer’s disease amyloid beta-peptide (Abeta40) in unilamellar liposomes is determined by anionic phospholipids, Peptides 27 (2006) 95–104.

[4] N. Arispe, H.B. Pollard, E. Rojas, The ability of amyloid beta-protein [A beta P (1-40)] to form Ca2+ channels provides a mechanism for neuronal death in Alzheimer’s disease, Annals of the New York Academy of Sciences 747 (1994) 256–266.

[5] N. Arispe, H.B. Pollard, E. Rojas, Giant multilevel cation channels formed by Alzheimer disease amyloid beta-protein [A beta P-(1-40)] in bilayer membranes, Proceedings of the National Academy of Sciences of the United States of America 90 (1993) 10573–10577.

[6] N. Arispe, E. Rojas, H.B. Pollard, Alzheimer disease amyloid beta protein forms calcium channels in bilayer membranes: blockade by tromethamine and aluminum, Proceedings of the National Academy of Sciences of the United States of America 90 (1993) 567–571.

[7] D.C. Bode, M.D. Baker, J.H. Viles, Ion Channel Formation by Amyloid-β42 Oligomers but Not Amyloid-β40 in Cellular Membranes, 292 (2017) 1404–1413.

[8] D. Cavallo-Medved, K. Moin, B. Sloane, Cathepsin B: Basis Sequence: Mouse, AFCS Nat Mol Pages 2011 (2011) A000508.

[9] E.N. Cline, M.A. Bicca, K.L. Viola, W.L. Klein, The Amyloid-β Oligomer Hypothesis: Beginning of the Third Decade, Journal of Alzheimer’s disease : JAD 64 (2018) S567–S610.

[10] S.R. Durell, H.R. Guy, N. Arispe, E. Rojas, H.B. Pollard, Theoretical models of the ion channel structure of amyloid beta-protein, Biophysical journal 67 (1994) 2137–2145.

[11] H.P. Erickson, Size and shape of protein molecules at the nanometer level determined by sedimentation, gel filtration, and electron microscopy, Biol Proced Online 11 (2009) 32–51.

[12] L. Fan, C. Mao, X. Hu, S. Zhang, Z. Yang, Z. Hu, H. Sun, Y. Fan, Y. Dong, J. Yang, C. Shi, Y. Xu, New Insights Into the Pathogenesis of Alzheimer’s Disease, Front. Neurol. 10 (2020).

[13] M.E. Guicciardi, M. Leist, G.J. Gores, Lysosomes in cell death, Oncogene 23 (2004) 2881–2890.

[14] C. He, D.J. Klionsky, Regulation mechanisms and signaling pathways of autophagy, Annual review of genetics 43 (2009) 67–93.

[15] B.L. Heckmann, B.J.W. Teubner, B. Tummers, E. Boada-Romero, L. Harris, M. Yang, C.S. Guy, S.S. Zakharenko, D.R. Green, LC3-Associated Endocytosis Facilitates β-Amyloid Clearance and Mitigates Neurodegeneration in Murine Alzheimer’s Disease, Cell 178 (2019) 536-551.e514.

[16] Y.-B. Hu, E.B. Dammer, R.-J. Ren, G. Wang, The endosomal-lysosomal system: from acidification and cargo sorting to neurodegeneration, Transl Neurodegener 4 (2015) 18–18.

[17] Z.S. Ji, R.D. Miranda, Y.M. Newhouse, K.H. Weisgraber, Y. Huang, R.W. Mahley, Apolipoprotein E4 potentiates amyloid beta peptide-induced lysosomal leakage and apoptosis in neuronal cells, The Journal of biological chemistry 277 (2002) 21821–21828.

[18] S. Jin, N. Kedia, E. Illes-Toth, I. Haralampiev, S. Prisner, A. Herrmann, E.E. Wanker, J. Bieschke, Amyloid-β(1-42) Aggregation Initiates Its Cellular Uptake and Cytotoxicity, The Journal of biological chemistry 291 (2016) 19590–19606.

[19] N. Kavčič, K. Pegan, B. Turk, Lysosomes in programmed cell death pathways: from initiators to amplifiers %J Biological Chemistry, 398 (2017) 289.

[20] S. Lee, Y. Sato, R.A. Nixon, Lysosomal proteolysis inhibition selectively disrupts axonal transport of degradative organelles and causes an Alzheimer’s-like axonal dystrophy, The Journal of neuroscience : the official journal of the Society for Neuroscience 31 (2011) 7817–7830.

[21] H. Lin, Y.J. Zhu, R. Lal, Amyloid beta protein (1-40) forms calcium-permeable, Zn2+-sensitive channel in reconstituted lipid vesicles, Biochemistry 38 (1999) 11189–11196.

[22] M.-c.A. Lin, B.L. Kagan, Electrophysiologic properties of channels induced by Abeta25-35 in planar lipid bilayers, Peptides 23 (2002) 1215–1228.

[23] T. Mirzabekov, M.C. Lin, W.L. Yuan, P.J. Marshall, M. Carman, K. Tomaselli, I. Lieberburg, B.L. Kagan, Channel formation in planar lipid bilayers by a neurotoxic fragment of the beta-amyloid peptide, Biochemical and biophysical research communications 202 (1994) 1142–1148.

[24] R.A. Nixon, J. Wegiel, A. Kumar, W.H. Yu, C. Peterhoff, A. Cataldo, A.M. Cuervo, Extensive involvement of autophagy in Alzheimer disease: an immuno-electron microscopy study, Journal of neuropathology and experimental neurology 64 (2005) 113–122.

[25] S.K. Rhee, A.P. Quist, R. Lal, Amyloid beta protein-(1-42) forms calcium-permeable, Zn2+-sensitive channel, The Journal of biological chemistry 273 (1998) 13379–13382.

[26] F.J. Sepulveda, J. Parodi, R.W. Peoples, C. Opazo, L.G. Aguayo, Synaptotoxicity of Alzheimer Beta Amyloid Can Be Explained by Its Membrane Perforating Property, PLoS ONE 5 (2010) e11820.

[27] D.B. Teplow, On the subject of rigor in the study of amyloid β-protein assembly, Alzheimers Res Ther 5 (2013) 39.

[28] B. Turk, J.G. Bieth, I. Björk, I. Dolenc, D. Turk, N. Cimerman, J. Kos, A. Colic, V. Stoka, V. Turk, Regulation of the activity of lysosomal cysteine proteinases by pH-induced inactivation and/or endogenous protein inhibitors, cystatins, Biological chemistry Hoppe-Seyler 376 (1995) 225–230.

[29] B. Turk, V. Stoka, J. Rozman-Pungercar, T. Cirman, G. Droga-Mazovec, K. Oresić, V. Turk, Apoptotic pathways: involvement of lysosomal proteases, Biological chemistry 383 (2002) 1035–1044.

[30] E. Wesén, G.D.M. Jeffries, M. Matson Dzebo, E.K. Esbjörner, Endocytic uptake of monomeric amyloid-β peptides is clathrin- and dynamin-independent and results in selective accumulation of Aβ(1-42) compared to Aβ(1-40), Scientific reports 7 (2017) 2021–2021.

[31] F.S. Wouters, G. Bunt, Cathepsin B pulls the emergency brake on cellular necrosis, Cell Death & Disease 7 (2016) e2170–e2170.

[32] A.J. Yang, D. Chandswangbhuvana, L. Margol, C.G. Glabe, Loss of endosomal/lysosomal membrane impermeability is an early event in amyloid Abeta1-42 pathogenesis, Journal of neuroscience research 52 (1998) 691–698.

[33] Z. Yin, C. Pascual, D.J. Klionsky, Autophagy: machinery and regulation, Microb Cell 3 (2016) 588–596.

[34] W.H. Yu, A.M. Cuervo, A. Kumar, C.M. Peterhoff, S.D. Schmidt, J.H. Lee, P.S. Mohan, M. Mercken, M.R. Farmery, L.O. Tjernberg, Y. Jiang, K. Duff, Y. Uchiyama, J. Naslund, P.M. Mathews, A.M. Cataldo, R.A. Nixon, Macroautophagy--a novel Beta-amyloid peptide-generating pathway activated in Alzheimer’s disease, J Cell Biol 171 (2005) 87–98.

[35] D.V. Zaretsky, M. Zaretskaia, Degradation Products of Amyloid Protein: Are They The Culprits?, Current Alzheimer research 17 (2020) 869–880.

[36] D.V. Zaretsky, M.V. Zaretskaia, Flow cytometry method to quantify the formation of beta-amyloid membrane ion channels, Biochimica et biophysica acta. Biomembranes (2020) 183506.

[37] D.V. Zaretsky, M.V. Zaretskaia, Mini-review: Amyloid degradation toxicity hypothesis of Alzheimer’s disease, Neurosci Lett 756 (2021) 135959.

